# Introduction of single nucleotide variations into the promoter region of the mouse kidney anion exchanger 1 gene downstream of a TATA box changes its transcriptional activity

**DOI:** 10.1101/198457

**Authors:** Takumi Takeuchi, Mami Hattori-Kato

## Abstract

(Introduction) Distal renal tubular acidosis (dRTA) is characterized by an impairment of urine acidification and can be caused by variations in genes functioning in α-intercalated cells such as anion exchanger 1 (AE1). AE1 encodes the Cl^-^/HCO3^-^ exchanger in erythrocytes (eAE1) and α-intercalated cells (kAE1). We previously reported that in human erythroid intron 3 containing the promoter region of human kAE1, a SNP (rs999716) 39 base pairs downstream of the TATA box showed a higher minor allele A frequency in incomplete dRTA patients and such a promoter region showed reduced activities, leading to the hypothesis that those with the A allele may express less kAE1, developing incomplete dRTA. Here, single nucleotide variations were introduced downstream of the TATA box in the murine erythroid intron 3 to investigate changes in the promoter activity for murine kAE1 mRNA. (Methods) The erythroid intron 3 of C57BL/6 was subcloned into the pGL4.17 reporter vector, leading to mu kAE1Pro-pGL4.17. Three types of G to A substitutions were introduced 33, 35, and 36 base pairs downstream of the TATA box in the murine kAE1 promoter region by inverse PCR (Var1, Var2, and Var3, respectively). The HEK 293 cells were transfected with vectors. After 24 hours, the firefly luciferase activity was determined. (Results and Discussion) The promoter activity of Var1, as well as that of Var2 to a lesser extent, was reduced compared with that of the wild-type. The introduction of variations such as Var1 and Var2 into the murine genome by genome editing may help to establish mouse models of incomplete dRTA.

## Introduction

An upper urinary tract stone is a very common disease, and the annual incidence is estimated to be 203.1 per 100,000 citizens in Japan ^1^. Various conditions including distal renal tubular acidosis (type 1 RTA, dRTA) can induce stone formation in the kidney ^2^. dRTA is characterized by an impairment of urine acidification due to the dysfunction of α-intercalated cells in the distal nephron ^3^. Urinary stones formed in dRTA patients usually include a calcium phosphate component. Potassium citrate therapy is useful for the prevention of calcium stone formation in acidotic patients ^4^.

Acquired dRTA can be observed in patients with a variety of systemic and urinary tract diseases ^2^. dRTA is also caused by variations in genes functioning in α-intercalated cells, i.e., cytosolic carbonic anhydrase 2 (CA2) ^6^, ATP6V1B1 ^8-10^ encoding the B1 subunit of H^+^-ATPase, ATP6V0A4 encoding the A4 subunit of H^+^-ATPase, and SLC4A1/AE1/Band3 ^11-13^ encoding the Cl /HCO3 - exchanger. The former three show autosomal recessive inheritance and the latter is autosomal dominant and autosomal recessive. Anion exchanger protein 1 (AE1) is a dimeric glycoprotein ^14^ with 14 transmembrane domains ^16^ and it participates in the regulation of the intracellular pH by Cl^-^/HCO3^-^ exchange across the cell membrane. In the kidney, it is located at the basolateral membrane of α-intercalated cells and functions in the secretion of H^+^ into the tubular lumen in cooperation with H^+^-ATPase and H^+^/K^+^-ATPase at the apical membrane ^3^.

Human SLC4A1/AE1/Band3 is a gene spanning 19.757 kb of genomic DNA situated on chromosome 17, q21-22. The gene consists of twenty exons and transcribes two kinds of mRNAs utilizing different promoters encoding the Cl-/HCO3- exchanger in erythrocytes (eAE1) and that expressed in α-intercalated cells in the kidney (kAE1). The promoter for human kAE1 is located in erythroid intron 3, and so the human kAE1 transcript lacks exons 1 through 3 of the eAE1 transcript ^18^.

We previously reported that in human erythroid intron 3 containing the promoter region of human kAE1, a single nucleotide polymorphism (SNP rs999716) showed a significantly higher minor allele A frequency in incomplete dRTA patients compared with those without dRTA ^19^. The promoter region of the kAE1 gene with the minor allele A at rs999716, 39 basepairs downstream of the TATA box, showed reduced promoter activities compared with that of the major allele G, leading to the hypothesis that patients with the A allele at rs999716 may express less kAE1 mRNA and protein in the α-intercalated cells, developing incomplete dRTA.

Similarly to human kAE1 mRNA, mouse kAE1 mRNA lacks sequences of exons 1, 2, and 3, resulting in an N-terminal truncation of 79 amino acids compared with the mouse eAE1 protein ^20-23^. The internal initiation site of this truncated transcript was later identified in intron 3 of the mouse eAE1 gene ^24^. Mouse kAE1 mRNA has an additional specific exon K1, which is located inside intron 3 and spliced to the 5-prime end of exon 4. A corresponding additional exon was also reported for human kAE1 mRNA ^25^.

We have introduced single nucleotide variations in the downstream sequence of the TATA box in intron 3 of the murine AE1 gene to examine if these variations change the promoter activity for murine kAE1 mRNA transcription, as was seen in the human kAE1 gene.

## Materials and Methods

Mouse kidney AE1 promoter region: Genomic DNA was extracted from the tail of the C57BL6 mouse using a kit (DNA Extractor® Kit, Wako, Osaka, Japan). Intron 3 of the mouse kAE1 gene containing a TATA box was amplified from 10 ng of the above genomic DNA by a pair of kAE1ProSen / kAE1ProAS primers (Table 1) in a 50-μL volume with 0.3 μM sense and antisense primers, 2 μL of MgSO_4_, 4% DMSO, and 1 unit of KOD plus polymerase as follows: 94°C for 2 min, 40 cycles (98°C for 10 sec, 60°C for 30 sec, and 68°C for 30 sec) of 3-step PCR. Sense and antisense primers were attached with SacI and HindIII restriction sites at the 5-prime ends, respectively.

**Table 1:**
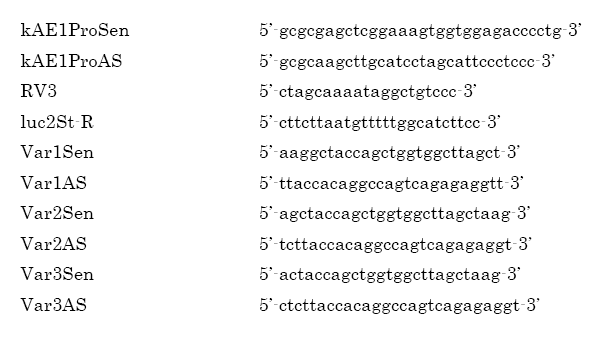
Amplification and sequencing primers.

PCR products were inserted using a DNA Ligation Kit (Mighty Mix, TaKaRa, Kusatsu, Japan) into the SacI / HindIII cloning site of pGL4.17 [luc2/Neo] reporter vector (Promega, Madison, WI, USA) following digestion with SacI and HindIII restriction enzymes, leading to the formation of a recombinant construct, mu kAE1Pro-pGL4.17. The DNA sequence of the construct was verified by Sanger sequencing using RV3 and luc2St-R primers (Table 1).

Variations introduced into the mouse kidney AE1 promoter region: In order to introduce three types of point variations downstream of the TATA box in the mouse kAE1 promoter region (intron 3 of the mouse AE1 gene), inverse PCRs were performed using three pairs of PCR primers (Var1Sen/Var1AS, Var2Sen/Var1AS, and Var3Sen/Var3AS) listed in Table 1 and 5 ng of mu kAE1Pro-pGL4.17 as a template in a 50-μL volume with 0.3 μM sense and antisense primers, 2 μL of MgSO_4_, 4% DMSO, and 1 unit of KOD plus polymerase as follows: 94°C for 2min, 10 cycles (98°C for 10 sec, and 68°C for 5 min) of 2-step PCR. Then, self-ligation of inverse PCR products was performed using a KOD-plus-mutagenesis kit (TOYOBO, Osaka, Japan). Competent cells (*E. coli* HST08, TaKaRa, Kusatsu, Japan) were transformed with those ligation products and vectors with variations in the mouse kidney AE1 promoter region were obtained and validated with Sanger sequencing using RV3 and luc2St-R primers. Three types of constructs with specific variations were named Var1, Var2, and Var3 (Figure 2).

**Figure 1.**
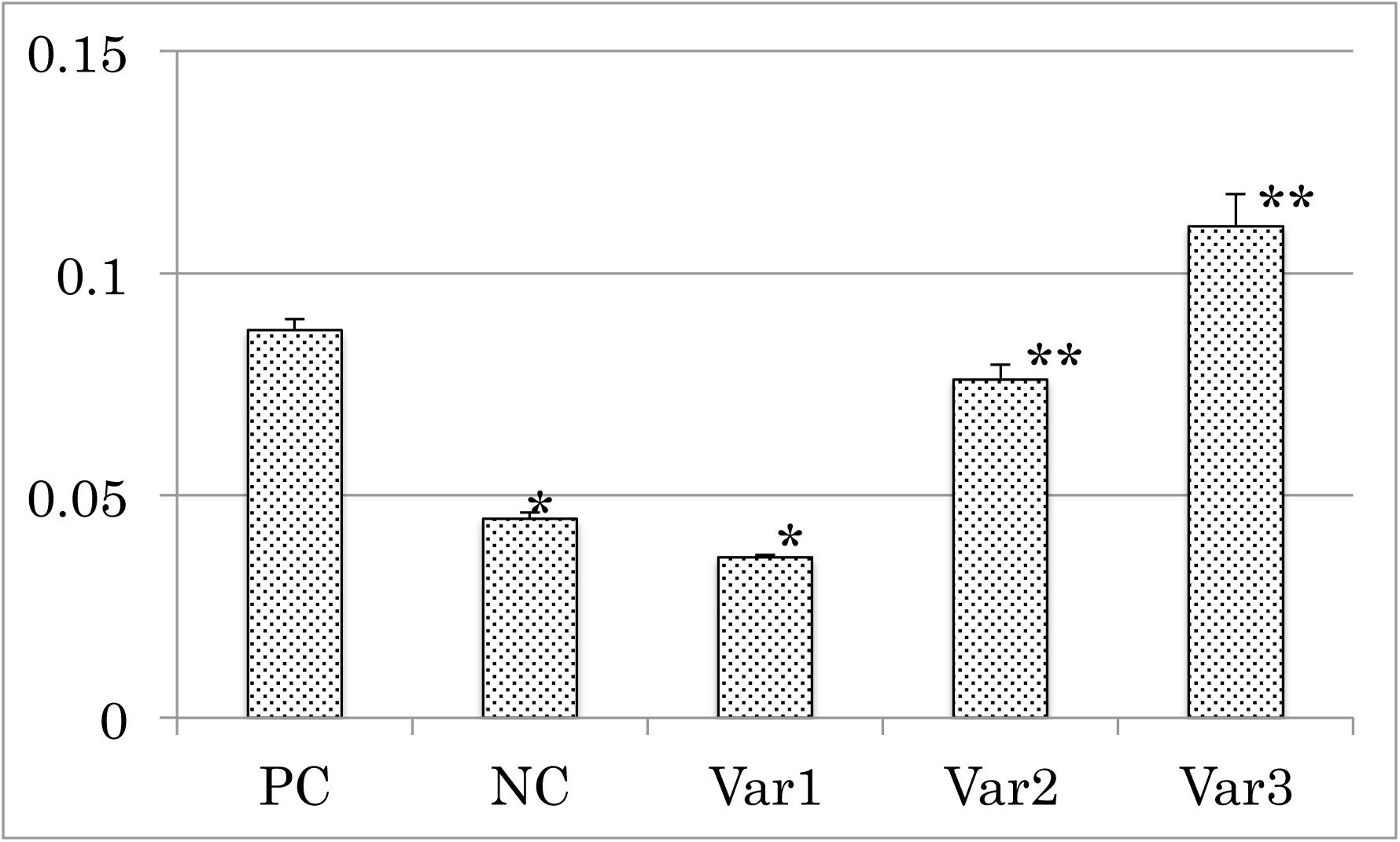
Promoter reporter luciferase assay using HEK293 cells. PC: positive control (mu kAE1Pro-pGL4.17, pGL4.17 containing the wild-type promoter for kAE1 mRNA), NC: negative control (pGL4.17), Var1, Var2, and Var3: G to A substitution 36, 38, and 39 base pairs downstream of the TATA box in mu kAE1Pro-pGL4.17, respectively, Vertical axis: the firefly luciferease activity divided by the internal control NanoLuc® luciferase activity, *: p<0.01 vs. PC, **: p<0.05 vs. PC by the t-test.

**Figure 2:**
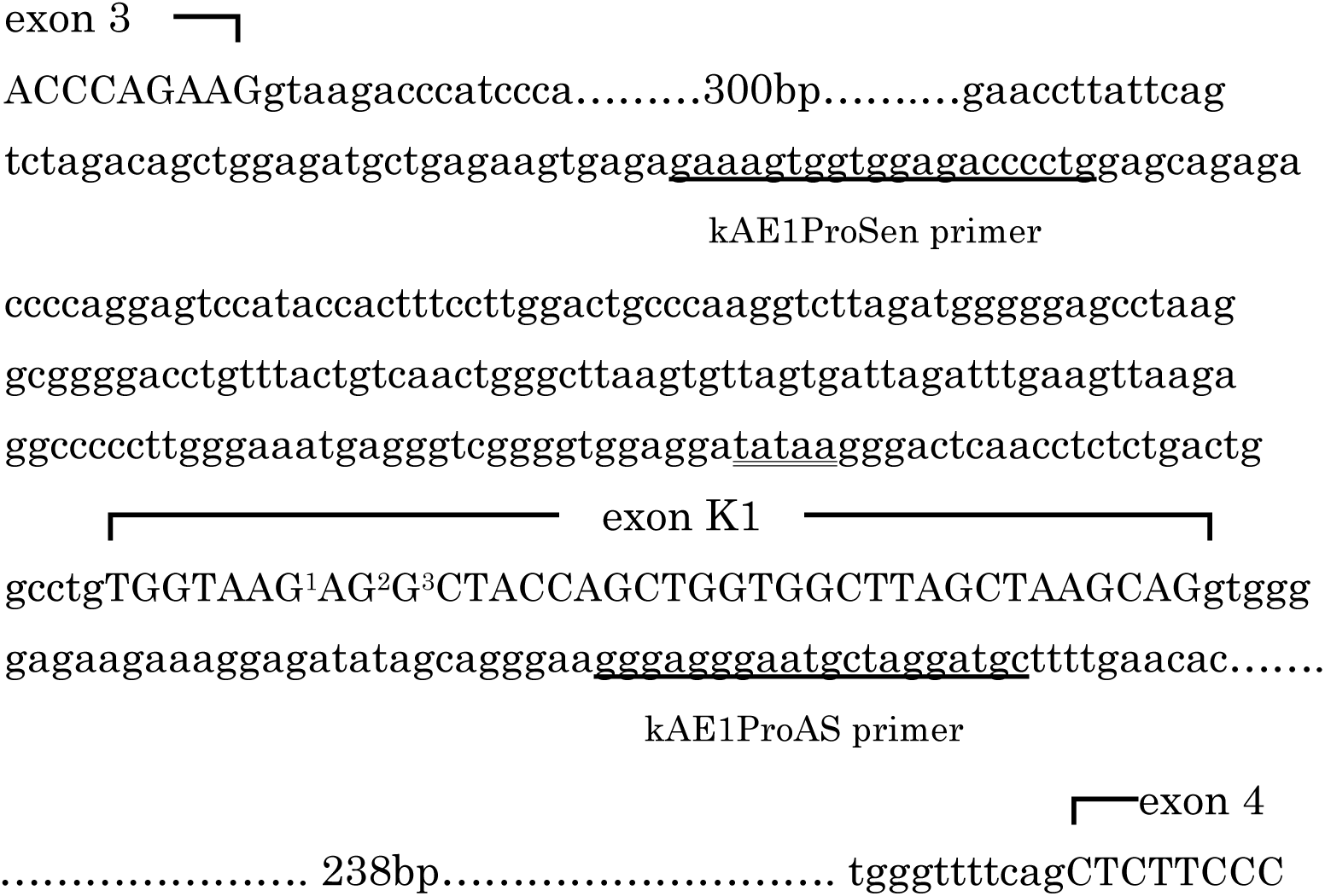
Intron 3 of the mouse AE1 gene. kAE1ProSen primer: sense amplification primer, kAE1ProAS primer: antisense amplification primer, exon K1: mouse kAE1 mRNA specific exon, K1, tataa: TATA box, G^1^, G^2^, G^3^: nucleotides where G to A substitutions were introduced (Var1, Var2, and Var3, respectively), Nucleotide numbering is according to Kopito et al.^20^

Promoter reporter assay: Human Embryonic Kidney 293 (HEK 293) cells (RBC 1637, Riken BRC, Tsukuba, Japan) were plated in a 96-well plate at a cell density of 2 × 10^4^ cells / well and were grown overnight in DMEM with 10% FBS. pGL4.17, Var1, Var2, Var3, and mu kAE1Pro-pGL4.17 vectors were used to transfect the HEK293 cells. The DNA mix for transfection was prepared in DMEM and consisted of 100 ng of the test plasmid and 1 ng of the pNL1.1.TK[Nluc/TK] control vector (Promega, Madison, WI, USA) expressing NanoLuc® luciferase under the control of the herpes simplex virus thymidine kinase promoter serving as an internal control to normalize firefly luciferase activity. The transfection was carried out using Lipofectamine 2000 (Invitrogen Corporation, Carlsbad, CA, USA). After 24 hours, the firefly luciferase activity was determined using the Nano-Glo^®^ Dual-Luciferase® Reporter (NanoDLR™) Assay System (Promega, Madison, WI) in a plate reader (Varioskan Flash, Thermo Fisher Scientific, Waltham, MA, USA). Each sample was quintuplicated. The normalized strength of the firefly luciferease activity was calculated by the firefly luciferease activity of each well divided by the NanoLuc® luciferase activity of the same well. For statistics, the t-test was applied.

## Results and Discussion

We have successfully developed three types of constructs with each specific variation in the promoter region of the murine kAE1 gene: Var1, Var2, and Var3. Variations were 33, 35, and 36 base pairs downstream of the TATA box, respectively. A G to A substitution was chosen for each variation, because a similar substitution was identified as a SNP in the human kAE1 promoter region downstream of the TATA box and it was strongly associated with type 2 renal tubular acidosis. Substitution sites were at the 7th, 9th, and 10th base pairs from the beginning of the transcription initiation site of kAE1 mRNA, inside the kAE1-specific exonK.

With the promoter reporter luciferase assay using HEK293 cells, the promoter activity of Var1, as well as Var2 to a lesser extent, was significantly reduced compared with that of the wild-type promoter region, as shown in Figure 1. On the contrary, the promoter activity of Var3 was enhanced compared with that of the wild-type. Considering that the SNP rs999716 located 39 base pairs downstream of the TATA box in the promoter region of the human kAE1 gene was associated with incomplete distal renal tubular acidosis, the introduction of point variations similar to Var1 and Var2 into the murine genome by appropriate means like genome editing may help establish mouse models of incomplete distal renal tubular acidosis. Such mice may show urolithiasis and mild osteoporosis as are seen in human incomplete distal renal tubular acidosis, without the more severe characteristics of complete distal renal tubular acidosis, such as growth retardation and fatal acidosis.

There are mouse models of dRTA. A mouse model that lacks AE1 (slc4a1-/-), but not heterozygous slc4a1+/- mice, exhibited complete dRTA characteristics such as spontaneous hyperchloremic metabolic acidosis, nephrocalcinosis, and hypocitraturia ^26^. Heterozygous and homozygous AE1 R607H knock-in mice corresponding to the most common dominant AE1 variation, R589H, causing dRTA in humans, displayed incomplete dRTA features ^27^. Those mice exhibit reduced basolateral AE1 and apical H^+^-ATPase expressions, while they show up-regulated Na^+^/HCO3^-^ co-transporters compared with the wild-type.

Na^+^/HCO3^-^ co-transporter knockout mice also had incomplete dRTA. Upon acid loading, those mice showed insufficient urinary acidification, hyperchloremic hypokalemic metabolic acidosis, and hypercalciuria with increased expression of the B1 subunit of H^+^-ATPase ^28^. Additionally, CA2-immunized mice with higher anti-CA2 antibody titers mimicking human Sjögren syndrome showed impaired urine acidification upon acid loading ^29^.

In conclusion, the engineered introduction of a G to A substitution into the 36 and 38 base pairs downstream of the TATA box in the murine kAE1 promoter region reduced the promoter activity, and may be a candidate method for establishing mouse models of incomplete dRTA and urolithiasis.

## Acknowledgments

This study was supported by a grant-in-aid for scientific research proposed by the Japan Labour Health and Welfare Organization. The Ethics Committee of Kanto Rosai Hospital approved the experiments. All experiments were performed in accordance with the relevant guidelines and regulations, including any relevant details.

